# Integration of brief light flashes varying in intensity and duration by the human circadian system

**DOI:** 10.1101/759134

**Authors:** Daniel S. Joyce, Manuel Spitschan, Jamie M. Zeitzer

## Abstract

The melanopsin-containing intrinsically photosensitive retinal ganglion cells (ipRGCs) are characterised by a delayed off-time following light offset. Here, we exploited this unusual physiologic property to characterise the exquisite sensitivity of the human circadian system to flashed light. In a 34-hour in-laboratory between-subjects design, we examined variable-intensity (3-9500 photopic lux; n=28 participants) full-field flashes at fixed duration (2 ms), and variable-duration (10 *μ*s-10 s) full-field flashes at fixed intensity (2000 photopic lux; n=31 participants) delivered using eye masks. We measured the circadian phase shift of the dim-light melatonin onset (DLMO) on the subsequent evening, acute melatonin suppression, objective alertness, and subjective sleepiness during the flash sequence. We find a clear dose-response relationship between flash intensity and the induced circadian phase shift, with an approximate increase of 10 minutes of phase delay for each ten-fold increase in photopic illuminance, but no parametric relationship between flash duration and induced circadian phase shift.

## Introduction

The human circadian system is exquisitely sensitive to light. Light exposure in the evening and night can acutely suppress the production of melatonin [1–6], shift the phase of the circadian clock [5, 7–11], and modulate alertness and vigilance [12–14]. This effect is mediated by the retinal photoreceptors, with a major role played by a subset (<3%) of the retinal ganglion cells that express the short-wavelength-sensitive photopigment melanopsin, rendering them intrinsically photosensitive (ipRGCs = intrinsically photosensitive retinal ganglion cells) [15]. ipRGCs also receive cone and rod input [16], which contribute to a complex signal driving the circadian system. The exact effect of a given light on the circadian system depends on its intensity, spectral distribution, duration, and circadian phase of administration [17–19]. While experimental durations of light exposure are typically on the order of hours, it has been shown that sequences of 2-millisecond flashes of bright light (~1,700 lux) can induce phase shifts in humans that are substantially larger than continuous light of the same illuminance [20].

Here, we systematically investigated the temporal integration properties of the human circadian system in a 34-hour in-laboratory between-subjects design. During the biological night, we exposed healthy observers (n=28) to a 60-minute sequence of short-duration white light flashes that varied in flash intensity over 4.5 orders of magnitude (3, 30, 95, 300, 950, 3000, or 9500 photopic lux) at fixed duration (2 ms) and measured the consequent impacts on circadian phase, melatonin suppression, and alertness. Additionally, we examined how short of a flash the human circadian system could respond to by examining sequences of short-duration light flashes spanning 6 orders of magnitude (10 *μ*s, 100 *μ*s, 1 ms, 10 ms, 100 ms, 1 sec, 10 sec) at fixed intensity (2000 lux). Stimuli were presented using eye masks, illuminating the retina with a homogenous full-field of light. Our results provide a mechanistic insight into the question of how the human circadian system integrates environmental information of ambient illumination.

## Results

### Circadian phase shifts to flashed lights are intensity-dependent and robust

We first examined our variable-duration data set for a dose-response relationship between the intensity of a white, broad-spectrum flash (CIE 1931 xy chromaticity: [0.4092, 0.3969], correlated colour temperature [CCT]: 3466K; melanopic efficacy of luminous radiation [ELR]: 0.72) measured in photopic illuminance, and the shift of the circadian clock measured as the difference in dim-light melatonin onset (DLMO) on subsequent evenings. Flash stimuli were delivered during the biological night as full-field homogenous stimuli using light masks and carefully calibrated in spectrum and temporal properties (Figures **S1** and **S2**). We find a significant effect of log_10_ illuminance (*F*(1, 25) = 5.68, *p* = 0.025, *R*^2^ = 0.185, *R*^2^_*adjusted*_ = 0.1524), indicating that ultra-brief (2 ms) flashes of light shift the circadian clock in an intensity-dependent manner (Figure **1*a***). Each increase of the illuminance by an order of magnitude while keeping the duration of the flash constant delays the circadian clock by approximately ten minutes (parameter estimate for log_10_ illuminance: *B*=0.17±0.070h).

**Figure 1.**
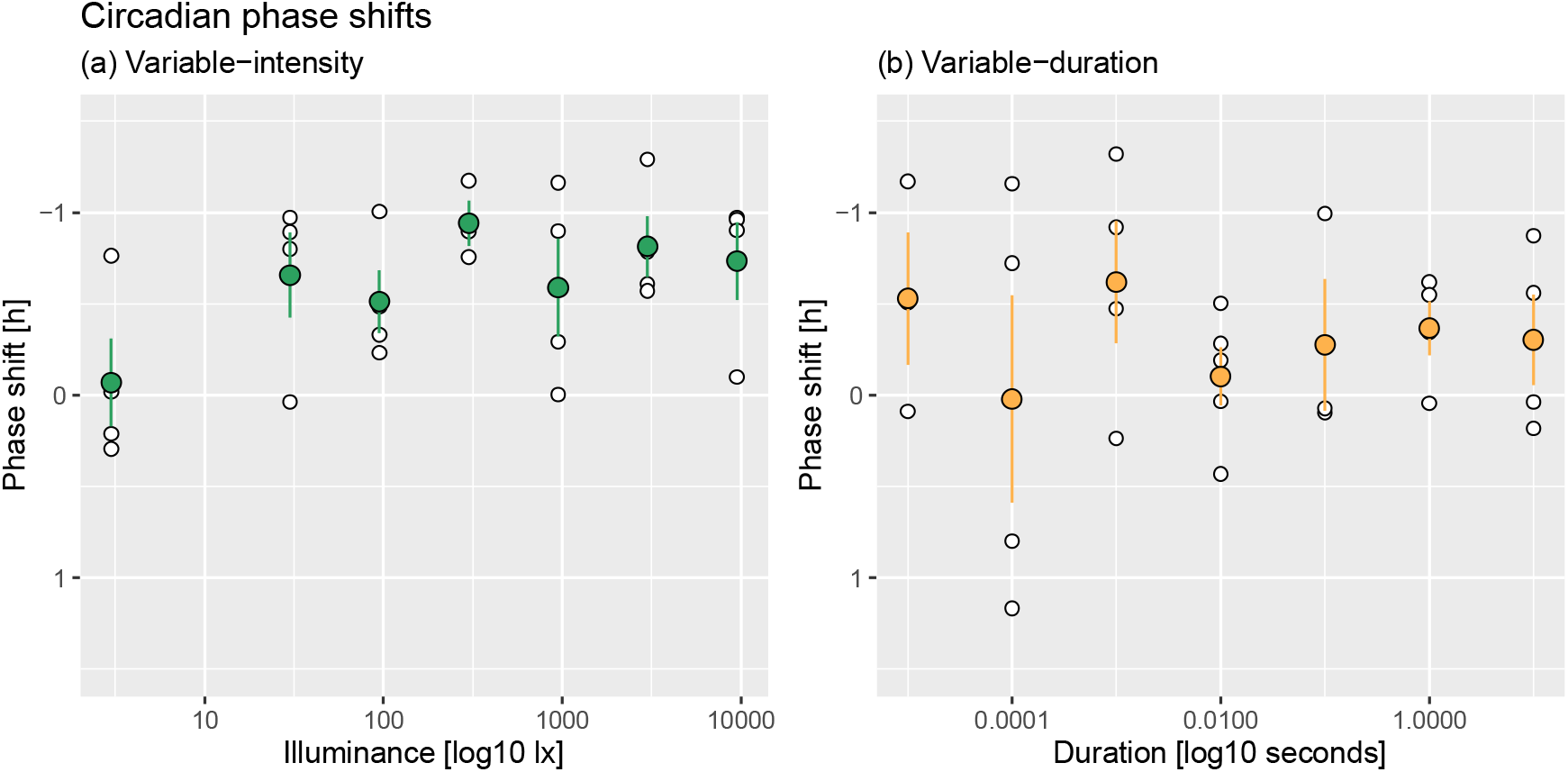
Flashes of light shift circadian phase in a illuminance-dependent manner. **a** Dose-response curve for circadian phase shifts across 3.5 orders of magnitude of photopic illuminance (3-9,500 lx; 2 ms flashes) measured in an in-laboratory between-subjects design (n=27). Individual, per-subject data points are shown as white circles, mean+SE estimates are shown as green circles. **b** Dose-response curve for circadian phase shifts across 6 orders of magnitude of flash duration (10 *μ*s-10 s; 2000 lx flashes) measured in an in-laboratory between-subjects design (n=27). Individual, per-subject data points are shown as white circles, mean+SE estimates are shown as orange circles.

We verified this effect by comparing this data to our previously published data on circadian phase shifts elicited in a parallel paradigm without any light stimulus [21]. We pooled the control data across two conditions in the original protocol corresponding to no administration of light while the participants were sleeping (n=7) or awake (n=7) (no difference in control phase shifts, Wilcoxon rank sum exact test *W*=21, *p*=0.71), and confirmed the findings by comparing the phase shifts elicited by our highest illuminance (9500 lux) with the phase shifts in the control condition (*W*=6, *p*=0.018). At this illuminance, a sequence of flashes elicits a phase delay of approximately 45 minutes (mean±SD phase delay=0.73±0.42). We verified that the phase angle of light onset was unrelated to the induced circadian phase shifts by examining the residuals of the linear model (Figure **S3;** R=−0.037, p=0.06 in variable-intensity protocol, R=0.14, p=0.5 in variable-duration protocol). In sum, these results clearly demonstrate that flashes induce a circadian phase shift graded by the illuminance of the flash.

We probed the shape of the dose-response relationship further by fitting logistic functions to the illuminance-phase shift data (Table **S2**). While individual data points are noisy, we find that a three-parameter logistic function best fits (as expressed using the Akaike Information Criterion, AIC) both mean and individual level data, with an estimated ED50 of 10.46 [-13.474, 34.395; 95% CI] lux (individual data fit) and 11 [−15.8441, 37.8491, 95% CI] lx (mean data fit), respectively. We note that the lower bounds of the estimate suggest negative illuminance, indicating overall noisy fits.

### No evidence for illuminance-graded acute effects of a sequence of ultra-brief light flashes on acute melatonin suppression, objective alertness and subjective sleepiness

We examined whether light flashes elicit acute changes in melatonin production, objective alertness (assessed using median reaction time measured using an auditory psychomotor vigilance test, PVT [22, 23]), and subjective sleepiness (assessed with the Stanford Sleepiness Scale [24]). We compared these endpoints just before administration of the light stimulus to the end of the hour of light administration. All effects are shown in Figure 2***a*, *c***, and ***e*** We find no effect of flash illuminance on acute melatonin suppression (*F*(1, 25) = 0.17, *p* = 0.68), objective alertness (*F*(1, 24) = 0.00084, *p* = 0.98), and subjective sleepiness (*F*(1, 25) = 0.86, *p* = 0.36). We find a significant decrease in subjective sleepiness independent of flash illuminance (mean±SD = −1.15±1.67; *t* = −2.37, *p* = 0.0261).

**Figure 2.**
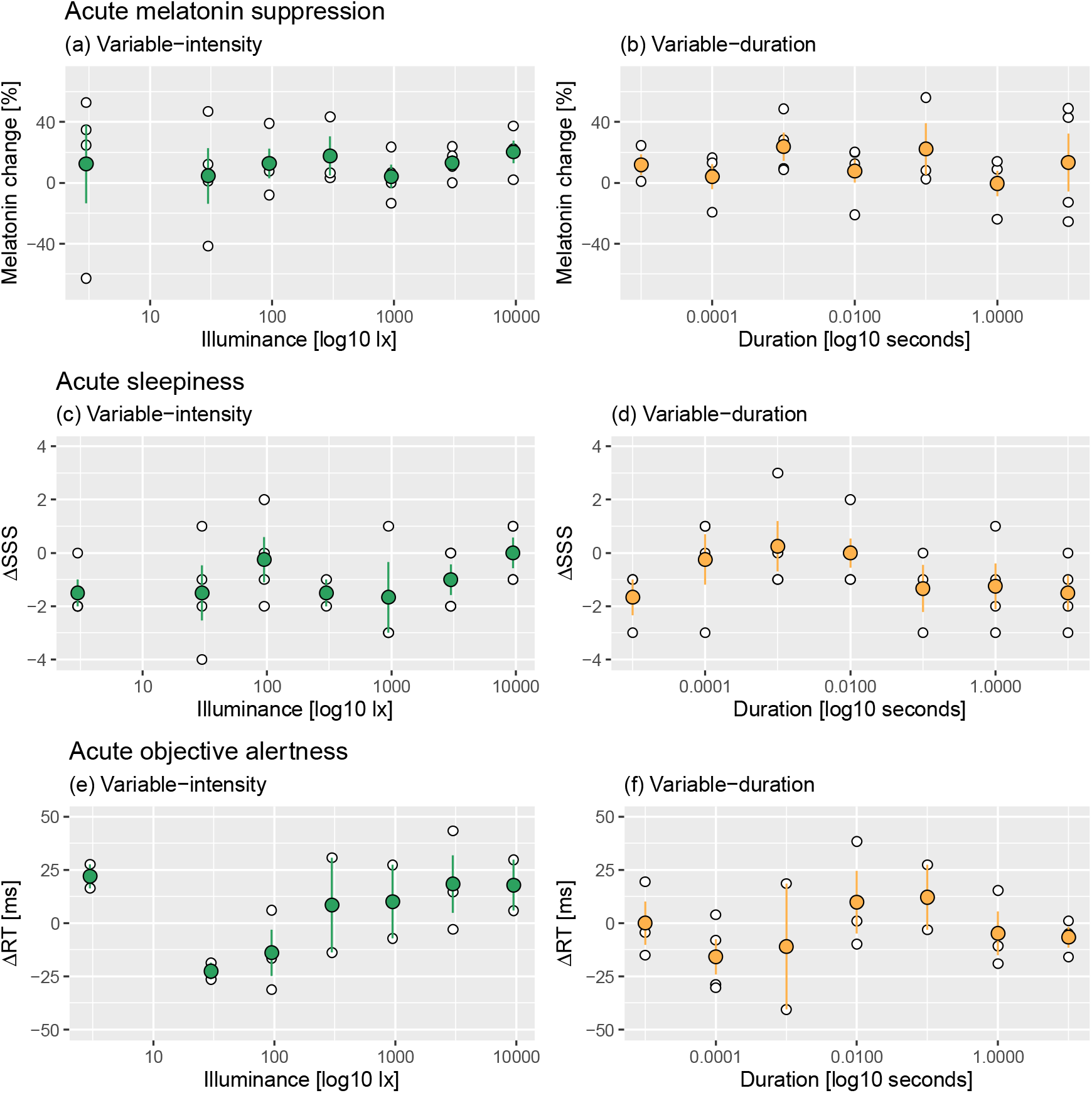
Flashes of light do not affect acute non-visual effects of light reliably. ***a, c, e*** Measurements of acute melatonin suppression, acute sleepiness and acute objective alertness in the variable-intensity protocol. **b, d, f** Measurements of acute melatonin suppression, acute sleepiness and acute objective alertness in the variable-duration protocol. Individual, per-subject data points are shown as white circles, mean+SE estimates are shown as green (variable-intensity; left column) and orange (variable-duration; right column) circles.

### Circadian phase shifts to flashed lights at fixed illuminance are duration-independent and non-robust

With flashes of varying intensity showing a clear intensity-dependent relationship with circadian phase shift, we examined the effect of a regime of 240 moderately bright flashes (~2,000 lux) of the same white, broad-spectrum light spectrum of varying duration (individual flash lengths of 10 *μ*s, 100 *μ*s, 1 ms, 10 ms, 100 ms, 1 sec, 10 sec; 15 second duration onset-to-onset). Across different flash durations, we find phase delays in circadian timing on the order of 18±36 minutes (Figure **1*b***). This effect is independent of the duration of flashes over the entire range of durations (*F*(1, 25) = 0.0018, *p* = 0.97), spanning six orders of magnitude (1:1,000,000). As with the variable-intensity protocol, we find no effect of flash duration on melatonin suppression (*F*(1, 25) = 0.023, *p* = 0.88), objective alertness (*F*(1, 25) = 0.11, *p* = 0.75), and subjective sleepiness (*F*(1, 25) = 0.85, *p* = 0.37). In parallel to our variable-duration results, we find a significant decrease in subjective sleepiness independent of flash duration (mean±SD = −0.741±1.56; *t* = −2.40, *p* = 0.0244).

## Discussion

As was implied in early studies of circadian photoreception [25], in response to brief flashes of light, the human circadian phase shifting system does not simply act as a photon counter, integrating over intensity and duration equally. As with long duration (6.5 hour) continuous light exposure presented during the early biological night [26], our data demonstrate a clear dose-response relationship between flash intensity and magnitude of the phase delay. Changes in the duration of the flash itself, however, seem to have little if any impact on strength of the light flash stimuli, as we observe invariant responses to a 1:1,000,000 difference in flash durations. Our data are consistent with light flashes being mediated through mechanisms that differ from those mediating responses to continuous light.

Much of the impact of continuous light on circadian function is thought to be mediated through the intrinsic (melanopsin) rather than extrinsic (rod/cone) photoreceptive circuits, especially for monochromatic light [27]. Our data, however, suggest a possible mechanism for temporal integration through a relative increase in the outer retinal rod and/or cone contributions to circadian photoreception vision via the extrinsic ipRGC pathway. In rodents, outer retinal inputs to the SCN are both measurable [28] and functionally significant in driving photoentrainment, as melanopsin-knockout mice can stably entrain to an LD cycle but exhibit attenuated circadian phase shifting to a single pulse of monochromatic blue light [29]. A subset of primate ipRGCs, the M1 subtype, receive excitatory input from the L and M cones, and inhibitory input from the S cones [30]. However, while there is converging evidence for connectivity of outer retinal inputs into ipRGCs, their consequence for circadian photoreception remains unclear. Cone signalling rapidly adapts under continuous light paradigms that greatly reduces its ability to signal for non-image-forming functions [31, 32]. For the circadian system at least, rod contributions may be preserved even at photopic irradiances and continue to drive photoentrainment [32, 33].

With our flashed light paradigm, the brief flashes in combination with the relatively long 15-second interstimulus intervals may allow rods to at least partially regenerate in the darkness [34, 35]. Because rods form the large majority of photoreceptors in the human retina, this contribution might not be insignificant. Future studies of *in vivo* ipRGC circuit electrophysiology, coupled with human studies using monochromatic light or photoreceptor-selective stimulation paradigms [36, 37] (rather than polychromatic, photoreceptor non-selective white light as used in this study) could clarify photoreceptor contributions to this process.

The ability of humans to consciously perceive light spans a very wide range of light intensities, from the sensitivity to single photons by the retinal rods [e.g. 38] to encoding fine spatial detail, colour and motion during daytime light levels. At detection threshold, the intensity and duration of a flash can be traded off against one another below a certain flash critical duration, leading to the same conscious perceptual performance if the product between the intensity and the duration is the same. For conscious perception, this temporal integration does not hold for flash durations over ~100 ms [39]. For shifting the circadian clock, however, it appears as though the mechanisms integrating light information are different and are likely non-linear, as shown in the results here.

We note that the variability of individual responses to flashed light is higher than that of continuous light, but commensurate with other studies of flashed lights in humans [21, 40] and with the recently identified large individual differences in circadian sensitivities to evening light exposure [41]. In comparison to continuous light paradigms, flashed light paradigms may be further be more susceptible to probabilistic photon catch over the short stimulus windows. The total stimulus duration over 1 hour is reduced 7,500-fold compared to an hour of continuous light or 48,750-fold compared to 6.5 hours of continuous light. In principle, differences in pupil size (which change the retinal illuminance) can modify melatonin suppression (at the same nominal corneal illuminance) [42]. However, the pupil only constricts to light onset with a delay of ~200 ms [43], making our variable-intensity flashes at 2 ms robust to such an effect. While there may be a cumulative effect of our sequence of flashes on long-term steady-state pupil size, the pupil will nonetheless reach significant redilation after our long 15 second inter-stimulus interval. While the variable-duration measurements at 1 and 10 seconds may be more susceptible to any such effect, we do not see a strong phase delay at the examined illuminance (2000 lux).

Flashed light confers advantages over continuous light when considering its acute effects on circadian physiology and behaviour. During the application of intermittent light, we did not find any significant dose-dependent effects on acute melatonin suppression, objective alertness and subjective sleepiness. This contrasts results for continuous light exposures during the biological light showing dose-responses relationships between light intensity and subjective sleepiness, EEG theta spectral power density [44], and melatonin suppression [26]. The lack of a dose-response relationship, and that light stimuli only altered subjective sleepiness, together suggest that these findings are underpinned by psychological factors (i.e., being awakened at night to observe flashed light) rather than psychophysical factors (differences in sensations of the light intensities). The distinct subtypes of ipRGCs may also contribute to this, where differences in retinal connectivity, temporal, spatial and intensity signalling, coupled with both distinct and overlapping brain targets [45–51], may lead to divergent light sensitivities depending on the physiologic pathway(s) assayed through the outcome measure(s) selected.

This study is the first to define the intensity sensitivity of the circadian system to sequenced flashes of light presented during the biological night. This paradigm leverages evolutionarily unusual stimuli to drive clinically meaningful shifts in circadian rhythms without substantial changes in acute measures of sleep behaviour and circadian physiology, in contrast to continuous light that often affects such performance markers in undesirable and disruptive ways. The flashed light paradigm is therefore a powerful method to drive clinically useful shifts in circadian rhythms, and, further, is orders of magnitude more efficient than continuous light paradigms in terms of time, energy and outcome, which is critical in the development of wearable technology that could be developed as a countermeasure to circadian desynchrony in a variety of environments.

## Materials and Methods

### Pre-registration and deviations from pre-registered protocol

The study protocol registered at ClinicalTrials.gov (NCT01119365; “Bright Light as a Countermeasure for Circadian Desynchrony”). The variable-duration study was pre-registered on the Open Science Framework (https://osf.io/5sv53/). Notably, we deviated from the pre-registered study protocol by including an additional (10 sec) exposure duration. In both the variable-intensity and the variable-duration studies, polysomnography (PSG) data were collected but not analysed.

### Ethical approval

The protocol was reviewed and approved by the Stanford University Institutional Review Board, conforming to the Declaration of Helsinki. Prior to any procedures, subjects signed informed consent forms.

### Sample characteristics

A total of 59 healthy, young (18-35 years) participants of normal weight with no somatic diseases, sleep disorders (Pittsburgh Sleep Quality Index, PSQI [52] ≤ 5), moderate chronotype (reduced Morningness-Eveningness Questionnaire, MEQ [53] ≥ 11 and ≤ 27), no history of substance abuse (Alcohol Use Disorders Identification Test, AUDIT [54] ≤ 7), no depressive symptoms (Center for Epidemiologic Studies Depression scale, CES-D [55] ≤ 17), no use of hormonal contraceptives (females only), and normal colour vision (assessed with Ishihara Plates [56]) completed the studies. Females attended the lab within four days after the onset of menses.

### Variable-intensity study

28 participants (14 female, 14 male) completed the study. We excluded one participant from further analyses due to mistiming of the light stimulus relative to their circadian phase, yielding a total sample of 27 subjects (n=27, mean±1SD age: 27±5.16) years. The break-down of intensity assignment was: 3 lux, 4 participants; 30 lux, 4 participants; 95 lux, 4 participants; 300 lux, 3 participants; 950 lux, 4 participants; 3000 lux, 4 participants; 9500 lux, 4 participants.

### Variable-duration study

31 participants (13 female, 18 male) completed the study. We excluded two participants (1 female, 1 male) because of contaminated melatonin assays, one participant due to mistiming of light (female), and one participant because of accidental light exposure in the morning (female), yielding a total sample of 27 subjects (n=27, mean±1SD age: 25.7±3.94 years). The break-down of duration assignment was: 10 *μ*s, 3 participants; 100 *μ*s, 4 participants; 1 ms, 4 participants; 10 ms, 5 participants; 100 ms, 3 participants; 1 s, 4 participants; 10 s, 4 participants.

### Design

Participants were exposed to a sequence of 240 light flashes of varying, logarithmically spaced intensity at fixed duration (2 ms flashes; 3, 30, 95, 300, 950, 3000, or 9500 photopic lux), or varying duration at fixed intensity (10 *μ*s, 100 *μ*s, 1 ms, 10 ms, 100 ms, 1 s, 10 s, 2000 lux) spaced 15 seconds apart (from onset to onset). Acute effects on melatonin suppression, objective alertness, subjective sleepiness, and electrophysiological correlates of arousal (polysomnography, PSG) were measured immediately before and at the end of the light exposure (LE). Effects of LE on circadian phase was measured as the change in melatonin onset determined on a constant posture protocol (CP1) prior to light exposure and a constant posture protocol the following day (CP2).

### Protocol

Participants took take part in a 16-day study protocol:

#### Days 1-14

Participants were instructed to maintain a regular sleep and wake time schedule at home (±30-minute window of bed time and wake time). Sleep-wake patterns were monitored using an actigraph (Actiwatch2, Philips, Bend OR) and a self-reported sleep diary [57]. From these data, the midpoint of sleep (MSP) was estimated and used as the midpoint of the in-laboratory sleep opportunity.

#### Day 15

The participant entered the laboratory during the late afternoon of Day 15. During the evening, the participant underwent the first constant posture procedure (CP1, 8-hour duration, beginning 8 hours before habitual bedtime). During this procedure, the participant was given isocaloric meals (Ensure, Abbott Laboratories, Chicago IL) every 60 minutes, yielding a total caloric intake matched to what they would have received during dinner (calculated using the Mifflin-St. Jeor formula; [58]). During CP1, objective alertness (auditory version of the Psychomotor Vigilance Task, PVT; [22, 23]) and subjective sleepiness (Stanford Sleepiness Scale, SSS; [24]) were measured every 60 minutes. Saliva was collected every 30 minutes in untreated polypropylene tubes. Four hours before MSP (typical bedtime), participants were given the opportunity to sleep in darkness. After 1 hour and 45 minutes of sleep time in darkness (2 hours and 15 minutes before MSP), participants were awakened in the dark. Saliva was collected and auditory PVT and SSS were administered in the dark. Starting two hours before MSP, participants were exposed to a 60-minute sequence of 240 full-field flashes through a custom-made mask for light delivery. At 20, 40 and 60 minutes into the light exposure, saliva was collected. During the last 10 minutes of light exposure, the auditory PVT was administered, followed by an SSS. The mask was then removed from the participant and the participant continued to sleep in darkness.

#### Day 16

The participant was awakened at their habitual wake time (4 hours after MSP) into a dimly light room (<10 lux) and received breakfast and lunch at usual times. During the evening, the participant had a second constant posture procedure (CP2, 10 hours in total, beginning 8 hours before habitual bedtime), during which the participant was given isocaloric meals every 60 minutes, yielding a total caloric intake matched to what they would have received during dinner (same number as on Day 15, but spread over 10 hours instead of 8). During CP2, objective alertness (auditory PVT) and subjective sleepiness (SSS) were measured every 60 minutes and saliva was collected every 30 minutes.

### Stimulus delivery

Binocular full-field flashes of differing durations (2 ms in variable-intensity study; 10 *μ*s, 100 *μ*s, 1 ms, 10 ms, 100 ms, 1 s, 10 s in variable-duration study) were delivered to the observer. A custom-light mask was constructed using modified welding goggles (Jackson WS-80 Shade 5.0 Cutting Goggles; Kimberly-Clark Professional, Mississauga, ON, Canada) containing an acrylic panel with three horizontally arranged LED strips (12 SMD LEDs each, Lumileds L235-4080AHLCBAAC0). The light from the LEDs was diffused using a piece of diffusing acrylic (TAP Plastics, Mountain View, CA). For additional diffusion, the participant wore ping pong ball halves cut out to match the shape of the eye’s orbit. The LEDs were pulsed using electronics developed in-house based on the Arduino Uno R3 microcontroller. In the variable-intensity study, we used we verified photopic illuminances at 3, 30, 95, 300, 950, 3000, or 9500 photopic lux as confirmed by a calibrated photometer (International Light Technologies ILT900, Peabody, MA, USA).

We verified the timing of our apparatus for the nominal flash durations 10 *μ*s, 100 *μ*s, 1 ms, 10 ms, 100 ms, and 1 s using an integrated photodiode and transimpedance amplifier (OPT101, Texas Instruments) connected to a digital oscilloscope (Tektronix TDS 2024C). We measured the logic-level control pulse sent from the microcontroller as well as the light output (Fig. S1). We averaged over 128 (10 *μ*s), 128 (100 *μs*), 128 (1 ms), 64 (10 ms), 16 (100 ms), and 16 (1 s) pulses. The maximum amplitude of the pulse is approximately constant across all nominal pulse durations, indicating that there is no shift in light intensity due to duration. Integrating the light output, the logarithm of the integrated light output over the pulse duration is linear with the logarithm of the nominal pulse duration. Spectral output was measured using a PR-670 spectroradiometer (Photo Research, Syracuse NY) (yielding calibrated radiance) for all stimulus durations, indicating stationarity of the spectrum across stimulus durations and across the entire stimulus protocol (240 flashes, 1h). The results of these validation measurements are shown in Figure S2.

### Spectral and α-opic properties of light

The spectrum of the light measured in the corneal plane corresponded to a warmish white light (CIE 1931 xy chromaticity 0.4092, 0.3969; correlated colour temperature [CCT] 3466K). The spectrum is visualised in Figure S1 and tabulated as a relative spectrum in Supplementary Table S1. Metrics related to the recent CIE S026/E:2018 [59] standard are given in Table 1 for unit illuminance (1 lux). The spectral invariance with stimulus duration is shown in Figure S2. The invariance of the spectrum means the α-opic irradiances simply scale proportionally at different illuminances.

**Table 1:** α-opic stimulus properties at unit photopic illuminance (1 lux) calculated using the free CIE S 026 α-opic Toolbox (v1.049, version dated 26 March 2020) implementing the CIE S 026/E:2018 standard [59]. To derive the α-opic irradiance and the α-opic equivalent daylight (D65) illuminance at other photopic illuminances, multiply the values by the photopic illuminance value. The α-opic efficacy of luminous radiation is a scale-invariant ratio that only depends on the relative spectrum.

|  | S-cone-opic | M-cone-opic | L-cone-opic | Rhodopic | Melanopic |
| --- | --- | --- | --- | --- | --- |
| **α-opic irradiance, [mW·m<sup>-2</sup>]** | 0.35 | 1.2 | 1.6 | 0.90 | **0.72** |
| **α-opic efficacy of luminous radiation, [mW·lm<sup>-2</sup>]** | 0.35 | 1.2 | 1.6 | 0.90 | **0.72** |
| **α-opic equivalent daylight (D65) illuminance [lx]** | 0.43 | 0.85 | 1.0 | 0.62 | **0.55** |

### Melatonin assay

Saliva (at least 1 mL) was collected using polypropylene tubes (Fisher Scientific, Hampton NH). Samples were centrifuged after collection, frozen at –20°C, then stored at –80°C within 24 hours. Salivary melatonin concentration was assayed according to the manufacturer’s instructions (Salivary melatonin ELISA #3402, Salimetrics, Carlsbad CA; assay range: 0.78-50 pg/mL, sensitivity = 1.37 pg/mL). For a given participant, all samples were assayed on the same plate.

### Objective alertness: auditory PVT

We used a modified auditory psychomotor vigilance test (PVT; [22, 23]) to measure objective alertness using a serial collection of simple reaction times to auditory stimuli generated by a piezo buzzer. The stimuli were spaced apart in time at random inter-stimulus intervals (ISIs) between 2 and 6 seconds (discrete steps: 2, 3, 4, 5, 6 second ISIs). Upon button press, the tone stopped and the next trial began with a random ISI. Approximately 100 of these stimuli were presented, with the order of ISIs randomized at the beginning of the experiment. This assessment took 10 minutes. The auditory PVT was implemented using custom-made Arduino hardware and software. To measure response latencies, we modified sample code from a report validating using the Arduino platform to measure reaction times [60]. There is a 30-second time out which is considered a lapse trial. If there is a response during the ISI period, this was counted as an error of commission and the counter was reset, starting a new trial period. The random seed for the ISIs is initialized by reading analogue voltage noise from an unconnected pin in the Arduino.

### Subjective alertness: SSS

Participants completed the Stanford Sleepiness Scale (SSS, [24]). The SSS is a single question assessment of current sleepiness that uses a 7-point Likert-like scale. Scale values range from 1 to 7, with higher values indicating greater subjective sleepiness.

### Determination of phase shift

Phase shifts were determined by examining the acute change in the timing of salivary dim-light melatonin onset (DLMO). This onset was determined by calculating the time at which the melatonin concentrations rose above a variable threshold (twice the average of the first three daytime samples [61]). In cases in which this variable threshold was ambiguous (n=1 variable-intensity; n=4 in variable-duration study; 5/59 participants in total = 8.4%), we used a the hockey-stick curve-fitting method [62]. Determination of ambiguity was made blind to the lighting parameters to which the participant was exposed. Phase shift was calculated as the DLMO on CP1 – DLMO on CP2, such that negative changes indicate a delay in timing.

### Statistical analysis

Data were analysed using simple intercept+slope linear models using the lm() function in R, and non-parametric Wilcoxon rank sum exact test implemented using wilcox.test(). All code implemented these analyses along with the data are included in the data set. Dose response curves were fit using the drc package [63]. All code and data to reproduce the statistical analysis are available in the supplementary material.

## Acknowledgements

This work was supported by United States Department of Defense (W81XWH-16-1-0223 to J.M.Z.). M.S. is supported by a Sir Henry Wellcome Fellowship (Wellcome Trust 204686/Z/16/Z) and a Junior Research Fellowship from Linacre College, University of Oxford. The authors would like to thank Yvonne Quevedo, Cheng-Ann Wang and Steven Lai for recruiting participants and conducting the experimental sessions, Chun-Ping (Phoebe) Liao for melatonin analysis, and the Stanford Product Realization Lab for assistance with developing the light delivery system.

## Author Contributions

**Conceptualization:** Daniel S. Joyce, Manuel Spitschan and Jamie M. Zeitzer.

**Formal Analysis:** Daniel S. Joyce, Manuel Spitschan and Jamie M. Zeitzer.

**Funding Acquisition:** Jamie M. Zeitzer.

**Investigation:** Daniel S. Joyce, Manuel Spitschan and Jamie M. Zeitzer.

**Methodology:** Daniel S. Joyce, Manuel Spitschan and Jamie M. Zeitzer.

**Project Administration:** Jamie M. Zeitzer.

**Resources:** Jamie M. Zeitzer.

**Software:** Daniel S. Joyce and Manuel Spitschan.

**Supervision:** Jamie M. Zeitzer.

**Visualization:** Daniel S. Joyce, Manuel Spitschan and Jamie M. Zeitzer.

**Writing – Original Draft Preparation:** Daniel S. Joyce, Manuel Spitschan and Jamie M. Zeitzer.

**Writing – Review & Editing:** Daniel S. Joyce, Manuel Spitschan and Jamie M. Zeitzer.

---

**Figure S1.**
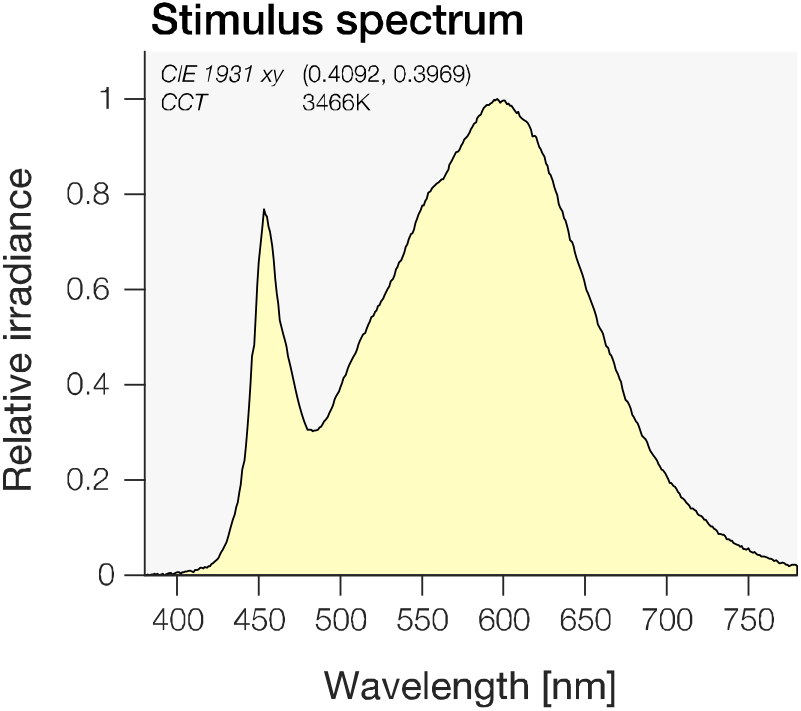
Stimulus spectrum (normalised to maximum irradiance). CCT was estimated using Psychtoolbox’s SPDToCCT function [64].

**Figure S2.**
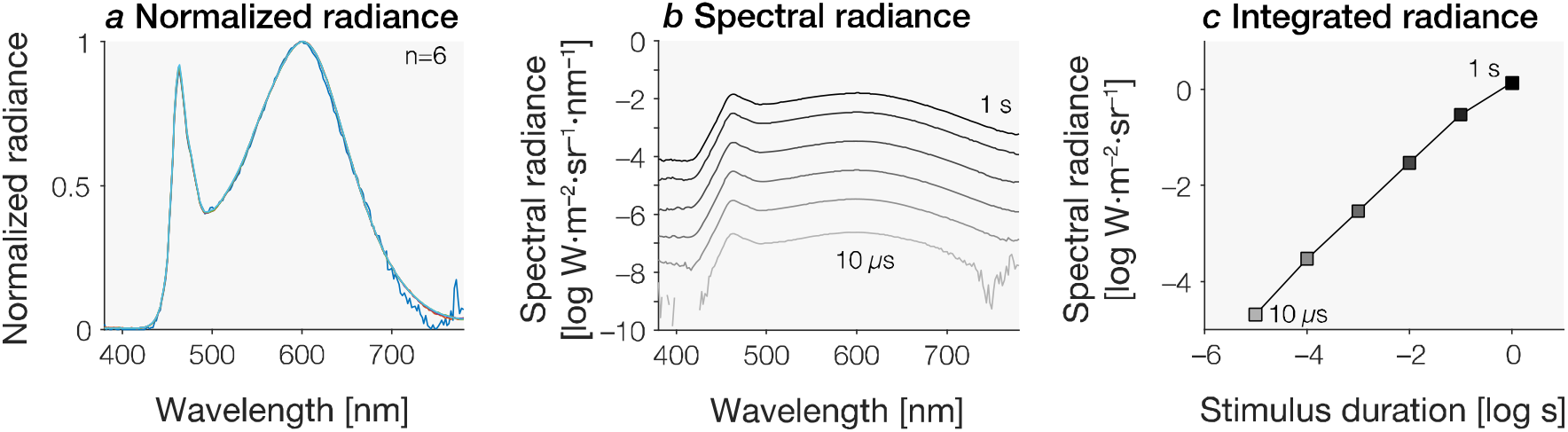
Spectral invariance across flash durations. ***a*** Time-averaged irradiance spectra measured across six flash durations (10 *μ*s-10 s, 2000 lx). The spectra are perfectly overlaying apart from long-wavelength noise for the lowest duration due to measurement noise at these low intensities. ***b*** Time-averaged irradiance spectra shown on a logarithmic scale. The noise in the short- and long-wavelength ends of the spectrum (>750 nm and <450 nm) appear amplified here but are inconsequential and represent amplified noise. The broken spectrum is due to the spectrometer reporting 0. ***c*** Time-averaged and spectrally integrated radiance, demonstrating linearity in spectrum across flash durations.

**Figure S3.**
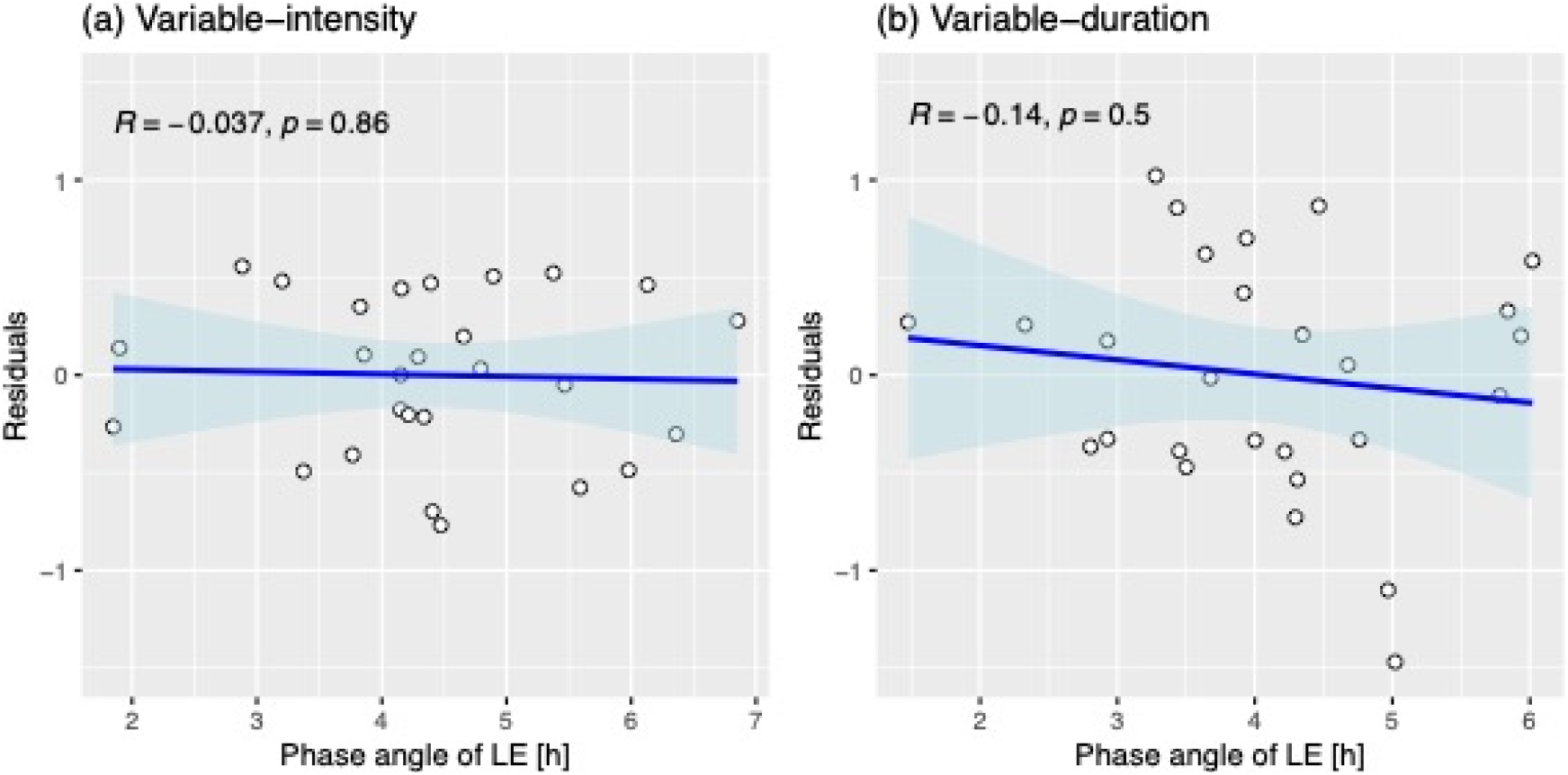
Relationship between phase angle of light exposure (LE) and residuals of linear model of phase shifts. There is no relationship between the timing of light exposure and the residuals in the linear model for induced phase shift in either the variable-intensity or variable-duration data.

**Table S2.**
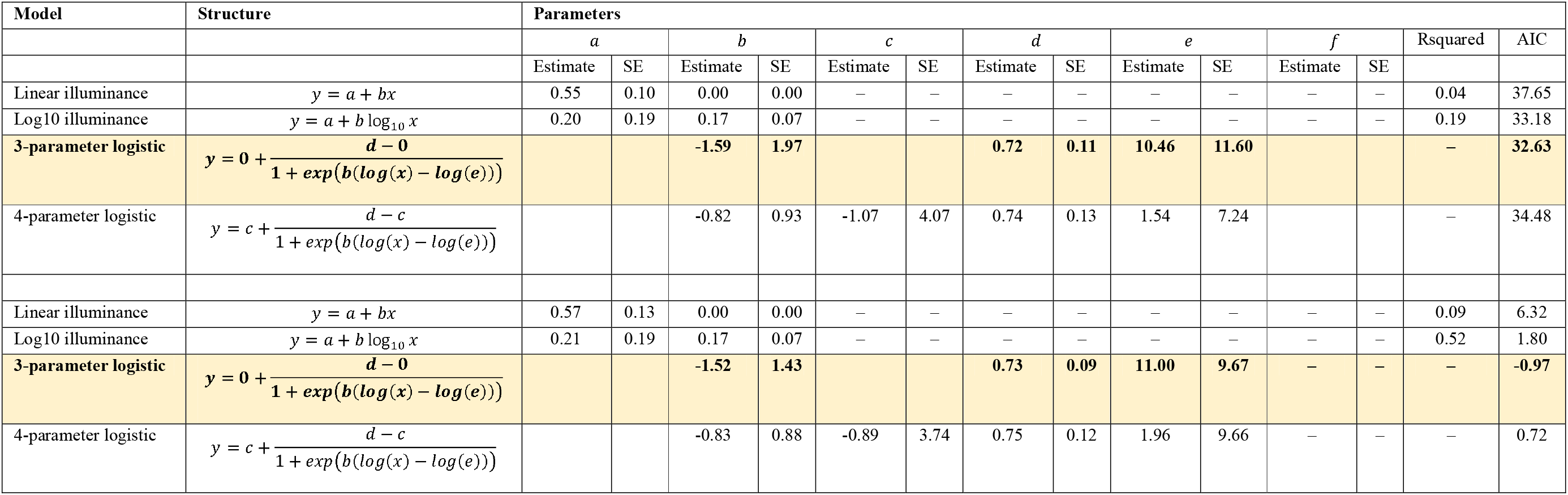
Model fits for the variable-intensity data using linear, log-linear and logistic functions with three, four and five parameters. We fitted both individual data, and per-illuminance average data. Goodness-of-fit is assessed using the Akaike Information Criterion (AIC). For model fits to both individual and mean data, a 3-parameter logistic function is the best fitting model according to the AIC (bold and highlighted in yellow). The meaning of parameters between the linear and log10 models, and the logistic models is not directly comparable.

